# Mechanical alterations of the hippocampus in the APP/PS1 Alzheimer’s disease mouse model

**DOI:** 10.1101/2020.12.29.424668

**Authors:** Nelda Antonovaite, Lianne A. Hulshof, Christiaan F.M. Huffels, Elly M. Hol, Wytse J. Wadman, Davide Iannuzzi

## Abstract

There is increasing evidence of altered tissue mechanics in neurodegeneration. However, due to difficulties in mechanical testing procedures and the complexity of the brain, there is still little consensus on the role of mechanics in the onset and progression of neurodegenerative diseases. In the case of Alzheimer’s disease (AD), magnetic resonance elastography (MRE) studies have indicated viscoelastic differences in the brain tissue of AD and healthy patients. However, there is a lack of viscoelastic data from contact mechanical testing at higher spatial resolution. Therefore, we report viscoelastic maps of the hippocampus obtained by a dynamic indentation on brain slices from the APP/PS1 mouse model where individual brain regions are resolved. A comparison of viscoelastic parameters shows that regions in the hippocampus of the APP/PS1 mice are significantly stiffer than wild-type (WT) mice and have increased viscous dissipation. Furthermore, indentation mapping at the cellular scale directly on the plaques and their surroundings did not show local alterations in stiffness although overall mechanical heterogeneity of the tissue was high (SD~40%). Therefore, reported mechanical alterations at a tissue scale indicates global remodeling of the brain tissue structure.

## 1. Introduction

Neurodegenerative diseases are difficult to research due to the complex biochemical processes and limited physical access to the brain. Neurodegenerative diseases are characterized by accumulation of aberrant proteins, reactive gliosis and neural cell death. The exact molecular mechanisms leading to the disease are often not know. In addition, in the past two decennia, several studies have indicated the relation between mechanical factors and brain functioning. For example, the mechanical sensitivity of neuronal and glial cells to the stiffness of environment has been demonstrated in cell culture experiments and *in vivo* by altered morphology, growth and other biochemical properties [1, 2, 3, 4, 5, 6, 7]. Moreover, in some neurodegenerative diseases, such as demyelinating disorders, the mechanical properties have been shown to change together with the structure thereby raising questions about the involvement of mechanobiological processes in disease progression [8, 9, 10].

Alzheimer’s disease is a neurodegenerative disease and the most common form of dementia in elderly. The exact molecular and cellular cause of dementia is still elusive, and effective drugs to halt or reverse the disease are still lacking. The pathology consists of the formation of extracellular plaques by the accumulation of amyloid *β* peptide (A*β*) and hyper-phosphorylated tau protein as intracellular neurofibrillary tangles (NFTs). Activated microglia and reactive astrocytes have been identified as the key players orchestrating chronic inflammatory response, linked to the severity of neuronal dysfunction in AD and are found accumulated around plaques similarly to glial scarring [11, 12]. The chronic neuroinflammation causes disease-related symptoms, such as loss of neurons and synapses ultimately leading to memory problems and dementia [13]. In addition to the underlying biological processes in AD, the investigation of mechanical properties received attention in recent years as a potential biomarker for early diagnosis, as shown by magnetic resonance elastography (MRE) studies where brain elasticity and viscosity decreased in AD human patients [14, 15, 16, 17, 18, 19], and as a novel drug target for tissue regeneration [**?** 20].

AD is often studied on mouse models carrying human transgenes with AD-linked mutations in amyloid precursor protein (APP) and Presenilin-1 (PSEN1). These mice exhibit extensive A*β* pathology in the hippocampus and cortex without tauopathy [13]. Although experiments on mouse brain tissue allow using contact mechanical testing methods, which is considered to be a gold standard, only two studies have reported Young’s modulus values of cortex [21] and hippocampus [22]. Therefore, more appropriate viscoelastic characterization of the hippocampus, where amyloid plaques are present, is needed. Although MRE can be used on humans as it noninvasively induces shear waves (<1 μm amplitude) from which one can calculate viscoelastic properties, the technique is limited to relatively high frequencies (30-900 Hz) and a low spatial resolution (~mm), thus, contact mechanical testing is needed to provide reliable mechanical data at higher resolution and lower-frequency spectrum to shed some light on the mechanobiology of AD.

In this study, we performed oscillatory indentation mapping of hippocampal subregions of APP/PS1 and wild type (WT) mice brain tissue slices where results of the latter have been already reported [23]. Both the storage modulus and damping factor of APP/PS1 mice were higher than WT in a region-dependent manner. The number of A*β* plaques and an increase in glial fibrillary acidic protein (GFAP) staining were observed by imaging (immuno)histochemical stained slices. None of the two correlated with viscoelastic parameters. Moreover, direct indentation mapping of the plaques at a single-cell scale did not show a mechanical contrast with the surrounding tissue, suggesting that the observed stiffening of APP/PS1 mouse brain is a tissue rather than cellular scale phenomena.

## 2. Methods

### 2.1. Sample preparation for indentation measurements

4 animals of 6 and 9 month-old (2 and 2 mice, respectively) C57BL6/Harlan wild type (WT) mice, and 6 animals of 6 and 9 month-old (5 and 1 mice, respectively) APPswe/PS1dE9 (APP/PS1) double-transgenic mice, which were littermates to WT mice, were used for indentation experiments reported in Section 3.1 [24]. All experiments were performed by following protocols and guidelines approved by the Institutional Animal Care and Use Committee (UvA-DEC) operating under standards set by EU Directive 2010/63/EU. 2 animals of 9-month-old APPswe/PS1dE9 double-transgenic mice were used for indentation experiments reported in Section 3.2. Animal handling and experimental procedures were previously approved by the Animal Use Ethics Committee of the Central Authority for Scientific Experiments on Animals of the Netherlands (CCD, approval protocol AVD1150020174314). Experiments were performed according to the Directive of the European Parliament and of the Council of the European Union of 22 September 2010 (2010/63/EU).

The mice were decapitated, dissected, and stored in an ice-cold carbonated 30% sucrose solution. Slices were cut in a horizontal plane with a thickness of approximately 300 μm using a VT1200S vibratome (Leica Biosystems, Germany) and placed to rest in artificial cerebrospinal fluid (aCSF) for 15 min in 32°C and 1 h at room temperature. Afterward, a single brain slice was placed in a perfusion chamber coated with 0.05% poly(ethyleneimine) solution, stabilized with a harp, and supplied with aCSF solution at 1 ml/min flow rate. Slices from 3 to 4 mm of dorsal-ventral positions were used in the experiment to minimize the effects of structural variation along with the hippocampus. Indentation measurements were performed within 8 h after extraction at room temperature. Results from WT mice brain slices were published previously (see [23]).

### 2.2. Dynamic indentation setup and measurement protocol on 300 μ m thickness brain slices

The setup and measurement protocol used in this experiment has been described previously [23]. In short, the custom indenter was mounted on top of an inverted microscope (Nikon TMD-Diaphot, Nikon Corporation, Japan) to image the slice during the measurements with a 2 × magnification objective (Nikon Plan 2X, Nikon Corporation, Japan) and a CCD camera (WAT-202B, Watec), while a cantilever-based ferrule-top force sensor equipped with spherical tip was indenting the brain slice from the top. Fig. 1 B shows a schematic drawing of the setup and a microscope image where the cantilever can be seen through the brain slice. The force sensors used for the experiments had cantilevers with the spring constant of 0.2-0.5 N/m and 60-105 μm bead radius. Indentation mapping was performed in indentation-depth controlled mode with the step size of 50-80 μm. Oscillations at 5.62 Hz frequency and 0.2 μm amplitude were superimposed on top of the loading ramp at an approximate strain rate of 0.01 *s^−^*^1^ (Fig. 1 A) to reach 8.5-15 μm indentation-depth (depending on the sphere radius), which corresponds to 7.5% strain 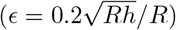 and fulfills small strain approximation *∊* < 8% [25]. Depth-dependent viscoelastic properties in terms of stor-age *E*′ and loss *E*″ moduli, and damping factor tan*δ* were calculated using dynamic mechanical analysis (DMA) which assumes linear viscoelastic mechanical behavior [26]:

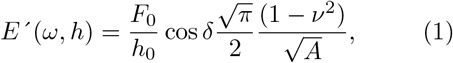

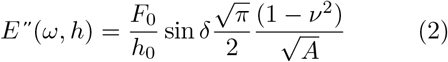

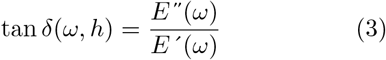

where *ω* is the oscillation frequency, *F*_0_ and *h*_0_ are the amplitudes of oscillatory load and indentation-depth, respectively, *δ* is the phase-shift between indentation and ol ad oscillations, *A* = *πa*^2^ is the contact area, 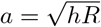 is the contact radius, *ν* is the Poisson’s ratio of compressibility (we assume that brain is incompressible *ν* = 0.5), *h* is the indentation-depth.

**Figure 1:**
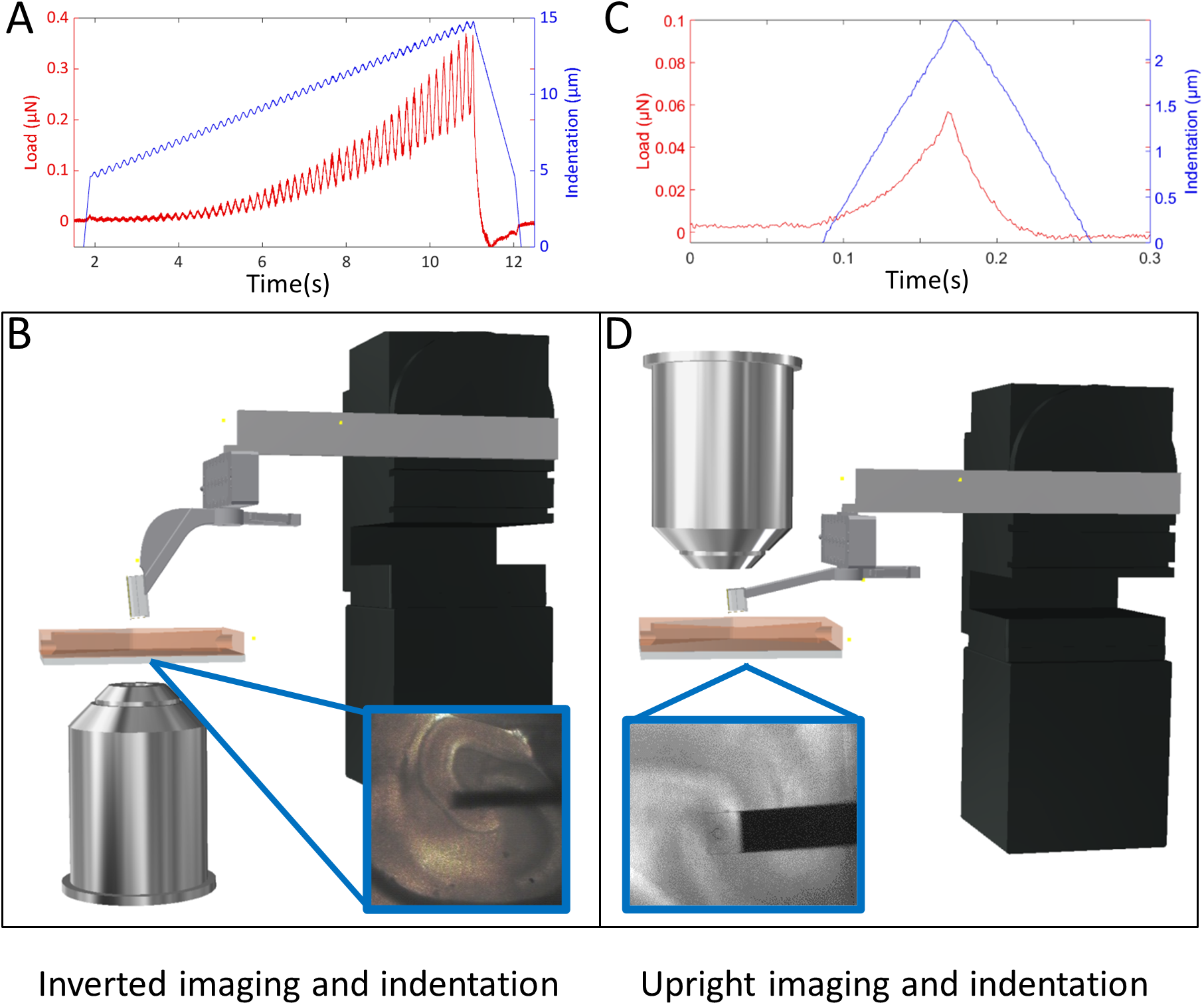
Indentation setup and profiles. The oscillatory-ramp indentation profile A) was used for viscoelastic characterization of the whole hippocampus at a tissue scale by using an inverted bright-field microscope and indenter B) to localize the sphere of the probe (R=60-105 μm) on top of the brain tissue. Static indentation profile C) was used to measure elastic properties at a cell scale with an upright fluorescence microscope and indenter D) by localizing the tip of the probe (R=21 μm) above the stained plaque.

Measurements were carried out on 6 slices from 6 APP/PS1 mice of which one was 9 month-old and the others were 6 month-old, with 63 – 535 measurement points per slice and 1235 total number of indentations. During the same experiment, indentation measurements (n=1029) were also performed on 5 slices from 4 WT mice (three 6-month-old and two 9-month-old) [23]. After the measurements, slices were stained with Methoxy X-04 (10 μM solution for 12 min), fixed in 4% paraformaldehyde (PFA) overnight at 4 °C, and imaged with a fluorescence microscope (Leica Microsystems, Wetzlar, Germany).

The second set of measurements was done with upright fluorescence microscope Axioskop 2 FS plus (Fig. 1 D), where the probe was modified to have a transparent cantilever tip, half-size ferrule, and shallow angle holder to fit the probe under the long working distance objective (Plan-Neofluar x5/0.16, Zeiss). An indentation probe with spring constant of 0.23 N/m and tip size of 21 μm was used to make static indentation mapping (see Fig. 1 C) with 5 – 9 μm step size at 30 μm/s piezo-transducer speed by indenting up to 3 μm indentation-depth (7.5% strain), which fulfills small strain approximation [25]. The Young’s modulus was obtained by fitting Hertz model [27]. APP/PS1 slices were live stained with Methoxy X-04 (10 μM solution for 12 min) before the measurements to locate A*β* plaques by fluorescence microscopy (filter set DAPI-50LP-A-000, Semrock). 4 slices from 2 animals (one 6-month-old and one 9-month-old) were used in these experiments to obtain 6 indentation maps (n=100 – 121 locations per map).

### 2.3. (Immuno)histochemistry of 30 μm thickness brain slices

(Immuno)histochemistry (Section 3.3) was performed on three 6 months old WT and three APP/PS1 female littermate mice from Jackson Laboratories [24]. Animal handling and experimental procedures were previously approved by the Animal Use Ethics Committee of the Central Authority for Scientific Experiments on Animals of the Netherlands (CCD, approval protocol AVD1150020174314). Experiments were performed according to the Directive of the European Parliament and of the Council of the European Union of 22 September 2010 (2010/63/EU). Mice were anesthetized with 0.1 ml Euthanimal 20% (Alfasan 10020 UDD) and transcardially perfused with 1X PBS. Brains were removed and collected in 4% paraformaldehyde for 48 hours before being transferred to 30% sucrose with sodium azide and stored at 4°C. Before cutting, brains were frozen in isopentane and mounted using Tissue-Tek (Sakura). Using a cryostat, brains were sliced horizontally in 30μm thick slices and collected in 1X PBS, which was then replaced by a cryopreservation medium (19% glucose, 37.5% ethylene glycol in 0.2 M PB with sodium azide) and stored at −20°C until further processing.

Slices were washed 3 times with PBS before they were blocked with 10% Normal Donkey Serum (NDS, Jackson ImmunoResearch, 017-000-121) and 0.4% Triton-X in 1X PBS for one hour at RT. Sections were incubated with different primary antibodies (Rat-anti-MBP, Sigma-Aldrich MAB386, 1:1000, monoclonal; Rabbit-anti-GFAP, CiteAb Z0334, 1:1000, polyclonal; Mouse-anti-6E10 Amyloid-*β*, BioLegend SIG-39300, 1:1000, monoclonal) diluted in 200 μl 10% NDS and 0.4% Triton-X blocking medium ON at 4°C. Thereafter, they were washed 3 times with 1X PBS and then incubated with 1:1000 secondary antibodies (Donkey-anti-Rat Cy3, Jackson ImmunoResearch 712-165-153; Donkey-anti-Mouse Alexa Fluor 488, Jackson ImmunoResearch 715-546-150; Donkey-anti-Mouse Alexa Fluor 594, Jackson ImmunoResearch 715-585-150; Donkey-anti-Rabbit Alexa Fluor 488, Jackson ImmunoResearch 711-545-152; Donkey-anti-Rabbit Alexa Fluor 594, Jackson ImmunoResearch 711-496-152) or 1:500 Wisteria floribunda agglutinin (WFA) dye diluted in 200 μl 3.3% NDS and 0.13% Triton-X in 1X PBS ON at 4°C, washed 3 times with 1x PBS and stained with 1:1000 Hoechst dissolved in 500μl 1x PBS for 10 min at RT. Slices were washed 2 times with 1X PBS and 1 time with MilliQ before mounting them on microscope slides using Mowiol (10% Mowiol (Millipore, 475904), 0.1% diazabicyclo(2,2,2)-octane, 0.1 M Tris and 25% glycerol in H_2_O; pH 8.5). Imaging was done with the Zeiss Axioscope.A1 epi-microscope operated with AxioVision software, using a 10x Plan-NeoFluar objective.

### 2.4. Statistics

Factorial (univariate) ANOVA analysis followed by post hoc tests with Bonferroni correction for multiple comparisons was applied during statistical analysis of data using IBM SPSS Statistics software.

## 3. Results

### 3.1. Viscoelastic properties of APP/PS1 mouse brain hippocampus is altered in comparison to WT

The depth-controlled oscillatory indentation mapping was performed at 50 – 80 μm axial resolution to capture regional mechanical differences hippocampus. The indentation lines were selected to cross the dentate gyrus (DG) and Cornu Ammonis 1 (CA1) or the CA3 fields of the hippocampus. As an example, Fig. 2 A shows indentation locations on the camera image of the APP/PS1 mice brain slice of DG and CA3 regions and storage modulus *E*′ and damping factor tan*δ* maps plotted at 7% strain (Fig. 2B and C, respectively). Different anatomical regions can be identified in the storage modulus map because of the high mechanical heterogeneity of different brain regions, which are related to varying morphology of brain regions (also observed on WT brain slices where data has been already published in [23, 28]); differences in damping factor tan*δ* are less pronounced but still visible.

**Figure 2:**
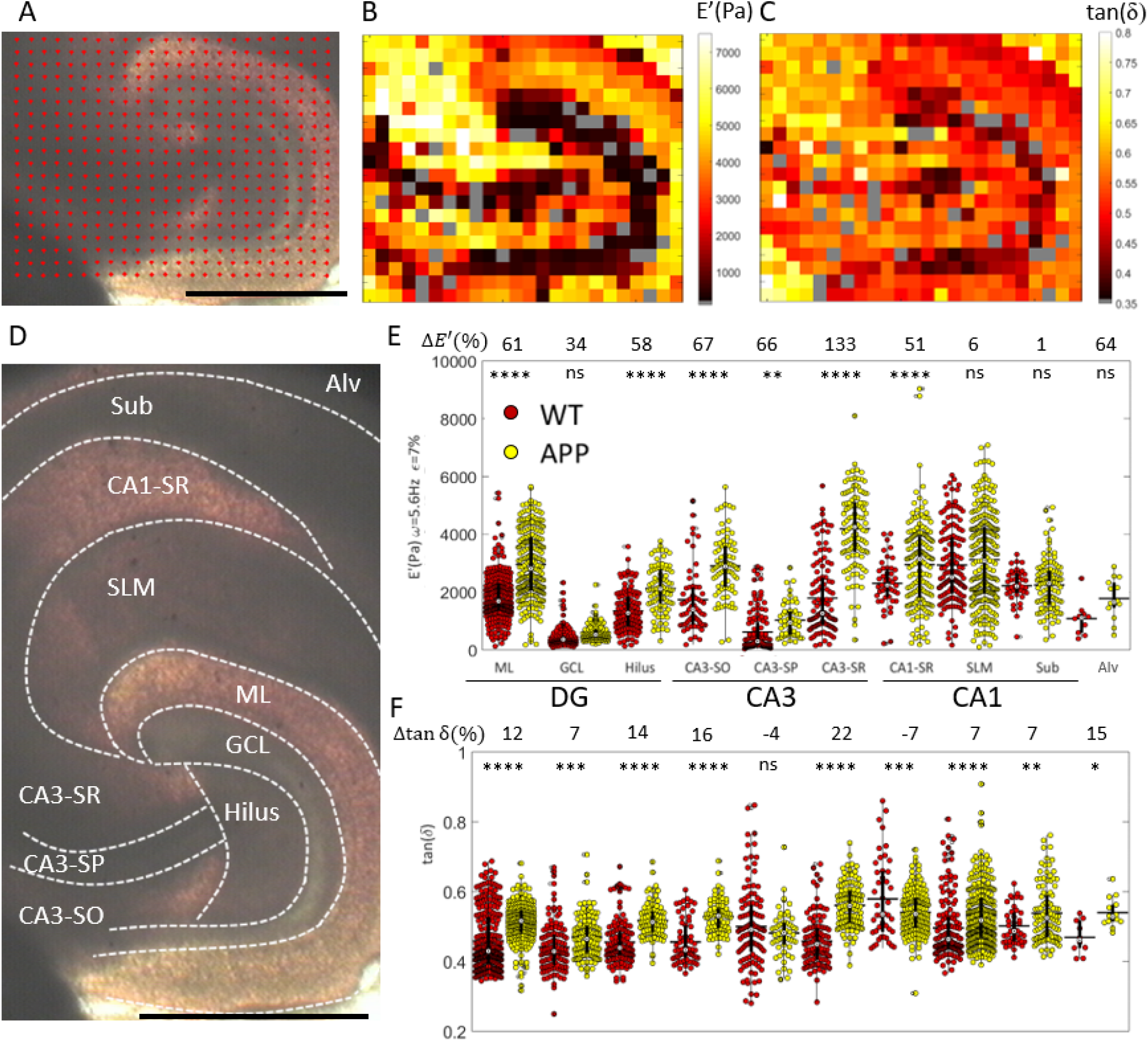
A) Red dots indicate indentation locations on a microscope image of the APP/PS1 mouse brain slice. B) Corresponding color map of storage modulus *E*′ and C) damping factor tan *δ* obtained at 5.6 Hz oscillation frequency and 7% strain. D) Hippocampal subregions were identified on the camera image of the slices with boundaries marked in dashed white lines. E) Storage modulus *E*′ and F) damping factor tan *δ* values of WT (red) and APP (yellow) mice hippocampal subregions at 7 0.5% strain. Data is merged over multiple slices (2 to 6 depending on the region, see Table S.1). The white dot indicates the median value with the vertical black bar for 25th and 75th percentiles and horizontal bar for the mean value. Bonferroni corrected p-values for pairwise comparison of simple main effects are indicated with asterisks:****p < 0.00001, ***p< 0.001,**p < 0.01, *p< 0.05, ns - non significant. Relative differences Δ(%) in estimated marginal means of storage modulus and damping factor are given above the graphs. Abbreviations: Alv - alveus, Sub - subiculum, SLM - stratum lacunosum moleculare, SR - stratum radiatum, SP - stratum pyramidale, SO - stratum oriens, ML - molecular layer, GCL - granule cell layer, dentate gyrus (DG) and cornus ammonis (CA1 and CA3). WT data has been reported previously [23].

Each indentation location was assigned to one of 10 measured hippocampal subregions (see Fig. 2 D) and storage modulus *E*′ and damping factor tan*δ* values at 7% strain are plotted in Fig. 2 E, F. 6 and 9 month-old data were merged together considering that both age groups are adults. The region, group of animals (WT or APP/PS1) and their interaction terms were significant factors for storage modulus and damping factor (p<0.0005, factorial ANOVA). The storage modulus of APP/PS1 mouse hippocampus was 1.5 times higher when considering all regions together with estimated marginal means of 1.63±0.05 kPa for WT and 2.39±0.04 kPa for APP/PS1 hippocampus. At an individual region level, the simple main effect analysis showed that differences in storage modulus were significant for the majority of the regions with the relative increase Δ*E*′ in these regions between 51 and 133% (Fig. 2 E). Moreover, the damping factor tan*δ* was 1.1 times higher for APP/PS1 than WT (0.522 0.003 and 0.481 0.003, respectively) when considering all regions together. At an individual region level, the damping factor was significantly higher for most of the regions with the relative increase Δtan*δ* between 7 and 22 %, except for CA3-SP, where the difference was not significant, and CA1-SR, where it was significantly lower (see Fig. 2 F). An increase in storage modulus *E*′ of APP/PS1 mouse hippocampus means that the material can resist more deformation while an increase in tan*δ* indicates that the loss modulus increased more than the storage modulus and, thus, the damping capability of the tissue, i.e. its fluid-like behavior, is larger.

In terms of mechanical heterogeneity, an elastic component (storage modulus) of mechanical behavior varies significantly between different regions, from 0.4 to 2.9 kPa for WT and from 0.6 to 4.2 kPa for APP/PS1 (estimated marginal means). Although damping is also different in different regions, it varies less, ranging between 0.44 and 0.58 for WT and between 0.47 and 0.56 for APP mouse hippocampal regions. Furthermore, the storage modulus *E*′ was obtained as a function of strain, between 5 and 7.5%, and showed a stiffening behavior with the strain *S* =Δ *E*′/Δ*ε* in the range of 0.1-1.1 kPa/% for WT mouse and 0.2 – 1.6 kPa/% for APP. The stiffening with the strain *S* was higher for all APP/PS1 hippocampal subregions except SLM. Interestingly, stiffer regions were stiffening more than softer regions (see Fig. S.1). To summarize, APP/PS1 mouse hippocampus shows a higher degree of mechanical heterogeneity in terms of storage modulus and a higher degree of nonlinearity with the strain.

Although plaques were present in these slices as confirmed by fluorescent staining with Methoxy X-04, mechanical heterogeneity of each region was high for both WT and APP/PS1 (mean SD~40% and 37%, respectively) and the resolution was low (50 – 80 μm) when compared to the size of the plaques (~10-50 μm diameter [29]). Furthermore, the staining and imaging of plaques were done after the indentation measurements, making identification of plaques in mechanical maps less accurate due to shrinkage of the slice during fixation with PFA. Taken together, while differences in viscoelastic parameters between APP/PS1 and WT hippocampal regions were assessed, it was not possible to identify the mechanical properties of individual plaques with this indentation protocol.

### 3.2. High resolution plaque mapping

To directly measure the mechanical properties of the plaques, the same side of the slice needs to be indented and imaged, thereby, the bright-field inverted microscope was replaced with an upright fluorescence microscope (see Fig. 1 D) and plaques were stained with Methoxy X-04 before starting the indentation measurements. The shallow-angle indentation probe was designed with a transparent cantilever to enable fluorescence imaging through the cantilever and a tip of R=21 μm was chosen to increase mapping resolution (5 – 9 μm). Fig. 3 shows Young’s modulus maps of the plaques and the surrounding areas. In non of the Young’s modulus maps, the plaques show strikingly different mechanical properties from the surroundings, although regional mechanical heterogeneity was high with a standard deviation between 39 and 52%.

**Figure 3:**
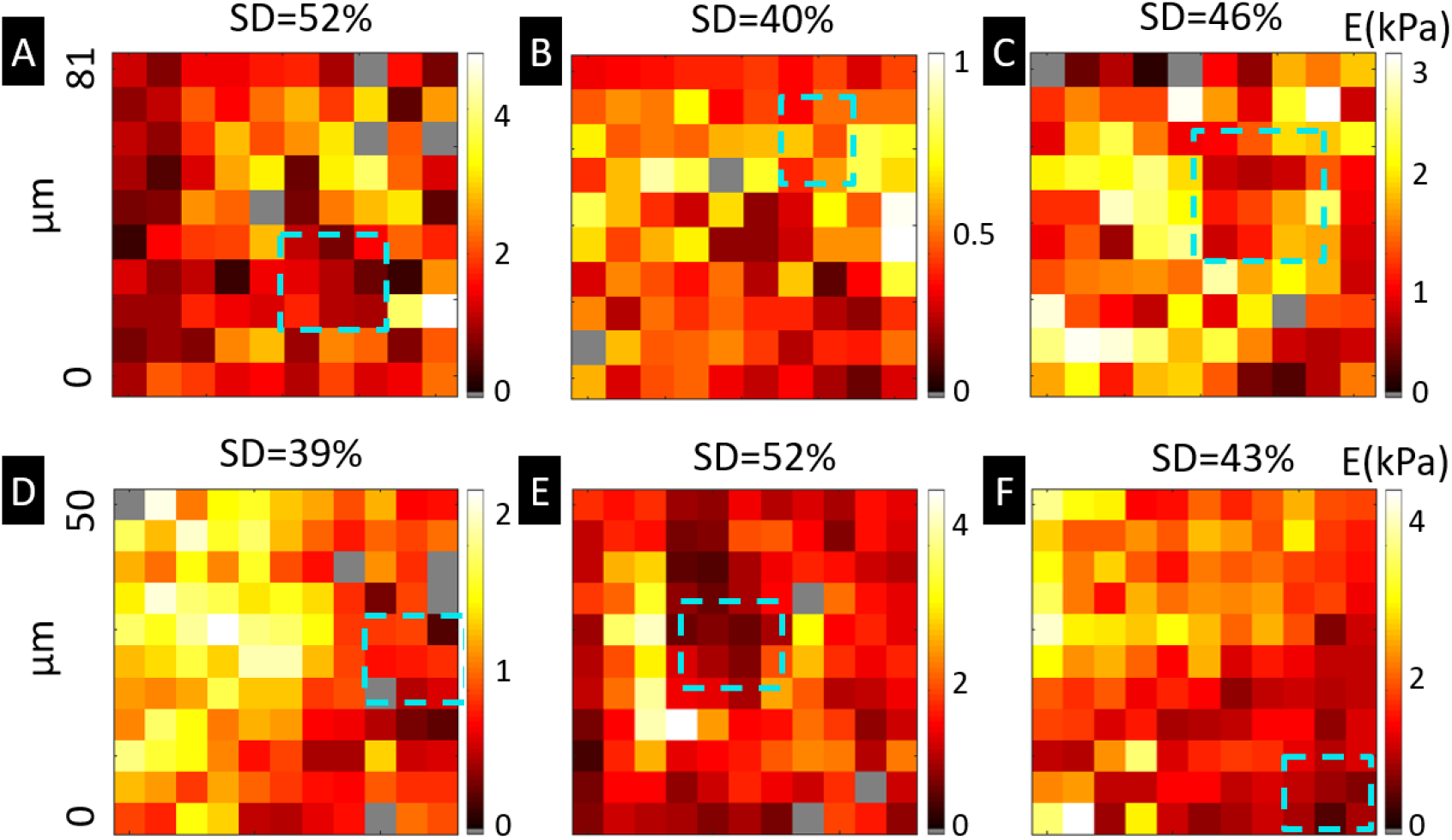
A-F) Maps of Young’s modulus obtained by fitting Hertz model up to 3 μm. The blue dashed line indicates location of the plaque obtained from staining with Methoxy-X04. A-C) Indentation mapping was performed on different slices from the same 6 month-old animal, regions A) ML, B) SLM/CA3-SR, C) SLM at 9 μm step size. D-F) Maps were obtained on one slice from 9 month-old animal, regions D) SLM, E) CA3-SO, F) Sub at 5 μm step size.

### 3.3. (Immuno)histochemical comparison

To investigate the underlying relationship between changes in mechanical properties and brain tissue composition in APP/PS1 mice, we performed (immuno)histochemical staining of the cytoskeleton of astrocytes (GFAP), cell nuclei (Hoechst), and A*β* plaques (6E10) (see Fig. 4 A). APP/PS1 mouse hippocampus have stained positive for the A*β* plaques with the mean number of plaques per region given in Fig. 4 B. SLM region had the highest averaged number of plaques - 4.3±2.6, followed by ML - 2.8±2.3 and Sub - 0.8±1, and all the other regions had rarely any plaques (<0.5). Furthermore, the plaques were surrounded by GFAP expressing astrocytes indicating astrogliosis. In comparison to mechanical results, SLM and Sub regions, did not show the difference in stiffness when comparing APP/PS1 and WT mice while ML region has increased stiffness by 1.6 times. Within the error of our measurements, there is thus no clear relationship between the number of plaques and stiffening. Additional WFA staining for perineural nets (PNNs) of extracellular matrix (ECM) and MBP for myelin did not show any qualitative differences in composition between APP and WT hippocampus (see Fig. S.2).

**Figure 4:**
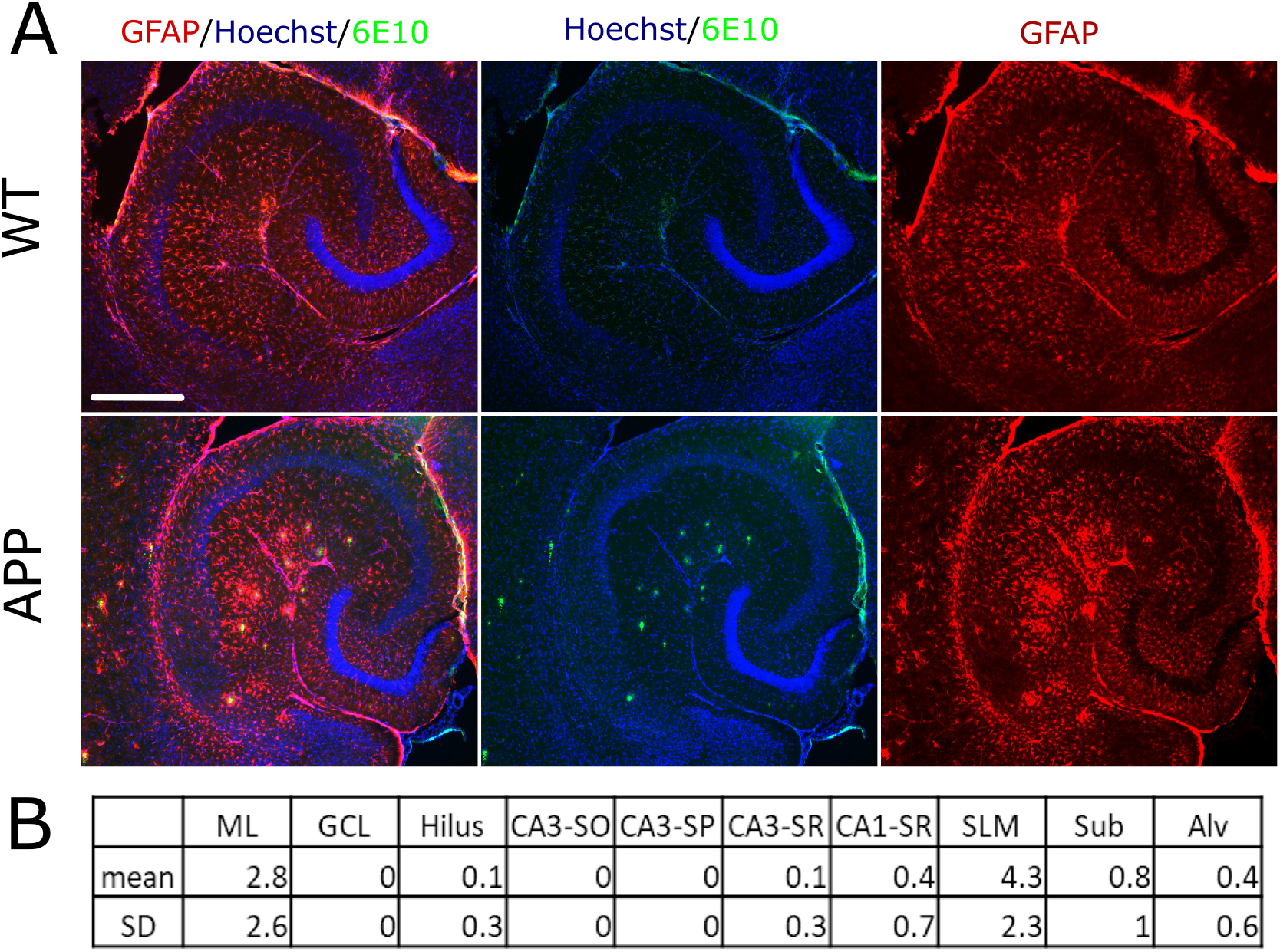
A) Fluorescent images of 6-month old WT and APP mice hippocampus stained with GFAP (astrocytes), Hoechst(nuclei) and 6E10 (A*β* plaques). The scale bar (white line) is 500 μm. B) Mean number of A*β* plaques per region (number of animals n=3, number of slices N=15).

## 4. Discussion

By using depth-controlled oscillatory indentation mapping we were able to obtain viscoelastic properties of 10 hippocampal subregions of APP/PS1 mice. We compared our results with data from WT mice, and found that the former is stiffer (e.i. higher storage modulus) and has better capability to dissipate mechanical energy (e.i. higher damping factor)(Fig. 2). We also showed that brain tissue from APP mice stiffens more with the strain and is more heterogeneous in terms of elasticity than WT mice (Fig. S.1). The number of plaques surrounded by upregulated GFAP astrocytes differs between the regions, but does not correlate with changes in viscoelastic parameters. Nevertheless, the increased viscoelasticity, both elastic resistance, and viscous dissipation, suggests that there are structural changes that take place during the development of plaques in the APP/PS1 mouse model that could affect the physiology of the brain. For example, one could speculate that similarly to how stiff brain implants induces an inflammatory response [3], brain stiffness change could contribute to the inflammatory response in Alzheimer’s disease.

It has been reported previously that APP/PS1 mouse brain has a decreased amount of myelin [21], an increased level of ECM proteins [30] and increased astrogliosis, marked by upregulation of GFAP expression surrounding plaques [31]. In this study, we performed (immuno)histochemical stainings (Fig. 4 and Fig. S.2) of cell nuclei (Hoechst), the cytoskeleton of astrocytes (GFAP), myelin (MBP), PNNs (WFA) and A*β* plaques (6E10) in an attempt to understand which structural components cause changes in viscoelastic parameters of APP/PS1 mouse hippocampus. We observed an increase in GFAP staining and the presence of A*β* plaques while there were no qualitative differences in other stained components. To better understand the structure-stiffness relationship, a more detailed quantitative study of brain tissue composition and organization is needed. The connection between brain compositions and its viscoelasticity is probably not a simple linear but a complex one with multiple interdependent variables such as alignment of fibers, density, size and type of cells, architecture and composition of extracellular matrix or fraction of water and lipids in the tissue. Furthermore, future studies should include multiple age groups from early to later stages of the disease to determine the onset of mechanical alterations.

Previously, MRE and atomic force microscopy (AFM) techniques have been used to compare healthy and AD mouse brains. AFM study on the fixated brain hippocampus reported lower Young’s modulus values for AD than healthy mice (40 and 104 kPa, respectively) although measurements were performed at the nm scale and the measured region was not identified [22]. Another AFM study has shown that Young’s modulus of fresh AD mouse brain tissue was lower than that of WT (0.4 and 0.7 kPa, respectively) when measured at 1.5 μm indentation-depth, although measurements were performed on cortex [21]. Unfortunately, the noise level of the indentation system used in this study was relatively high due to the perfusion flow, thus, it was not possible to fit the data below 1 μm indentation-depth. MRE studies on mice, similarly to humans (described in the introduction), have demonstrated a decrease in stiffness of AD mouse model brains in comparison to WT [32, 33], although the amount of deformation in terms of shear wave amplitude is at the μm scale and axial resolution is at ~mm scale. Our results are obtained by inducing deformation at a tissue scale e.i. 4-15 μm indentation-depth, and spatially resolving individual brain regions at ~50 μm axial resolution. Therefore, the difference in the mechanical deformation scale could explain the discrepancy between the studies.

High-resolution indentation mapping was done by modifying the setup to the upright fluorescence imaging configuration to localize and directly indent on the plaques. However, the mechanical heterogeneity of the regions was high (SD~40%) and the mechanical properties of individual plaques were not distinguishable from the surroundings (Fig. 3). There are two main limitations of the experimental approach used in this study: 1) imaging with fluorescence microscopy does not indicate how deep the plaque is situated within the brain tissue thickness; 2) the surface of the brain slice is rough and damaged due to slicing procedure, which results in an error in estimating mechanical properties, especially at smaller indentation-depths. Therefore, future experiments should include three-dimensional imaging, such as confocal microscopy [34, 35], to visualize how different structural components of the brain such as cells, ECM, neuronal projections, and plaques deform under compression and, thereby, extract their individual contribution to viscoelasticity.

## Code availability

The computer code used to generate the results of this study is available on request from the corresponding author.

## Data availability

All raw and processed data that support the findings of this study are available from the corresponding author upon request.

## Acknowledgments

The research leading to these results has received funding from the European Research Council under the European Union’s Seventh Framework Programme (FP/2007–2013)/ERC grant agreement no. [615170] and the Alzheimer Society in the Netherlands (Alzheimer Nederland WE.03-2017-04). The authors further thank M.Marrese for fruitful discussions, E.Paardekam for manufacturing indentation probes, and T.Smit for providing brain tissue slices and support in the lab.

## Author contributions

N.A., W.J.W., E.M.H., and D.I. designed the research; N.A. performed the experiments, analyzed the data, and wrote the manuscript; C.F.M.H. provided slices for high-resolution mapping; L.A.H. performed immunohistochemical stainings; all authors contributed to the editing of the paper.

## Supplementary Material

**Table S.1:** The number of slices used for testing and the total number of indentation points per region.

|  | Regions | ML | GCL | Hilus | CA3-SO | CA3-SP | CA3-SR | CA1-SR | SLM | Sub | Alv |
| --- | --- | --- | --- | --- | --- | --- | --- | --- | --- | --- | --- |
| WT | Number of slices | 5 | 5 | 5 | 5 | 4 | 4 | 4 | 5 | 3 | 2 |
|  | Number of indentations | 263 | 123 | 120 | 63 | 114 | 115 | 44 | 138 | 40 | 10 |
| APP | Number of slices | 4 | 4 | 4 | 3 | 3 | 3 | 6 | 6 | 5 | 2 |
|  | Number of indentations | 238 | 122 | 89 | 80 | 54 | 113 | 167 | 270 | 102 | 14 |

**Figure S.1:**
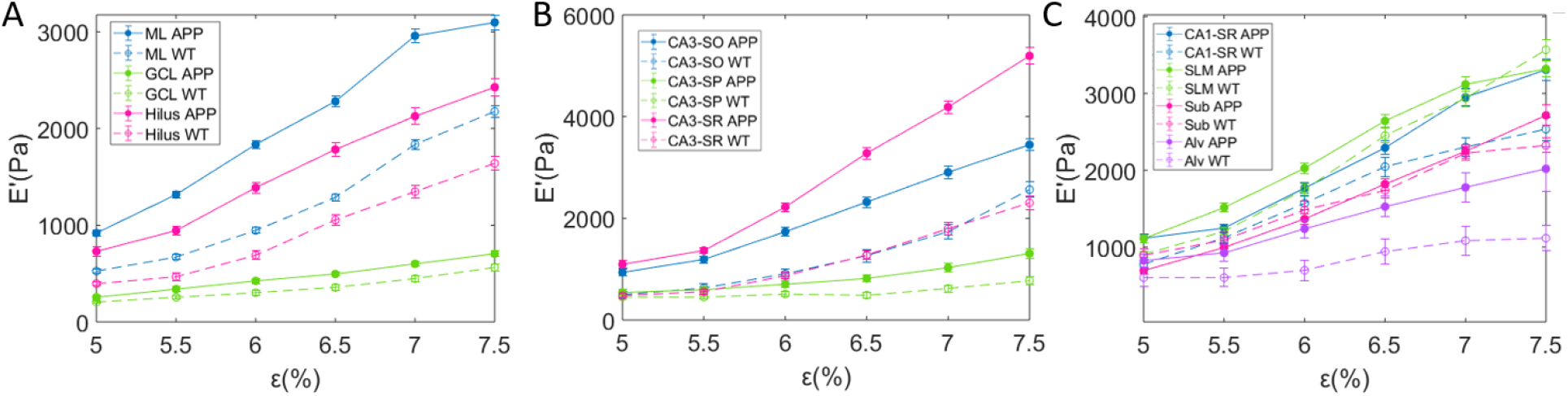
The mean storage modulus *E*′ as function of strain *ε* for different hippocampal regions of APP and WT mice. Error bars are SEM.

**Figure S.2:**
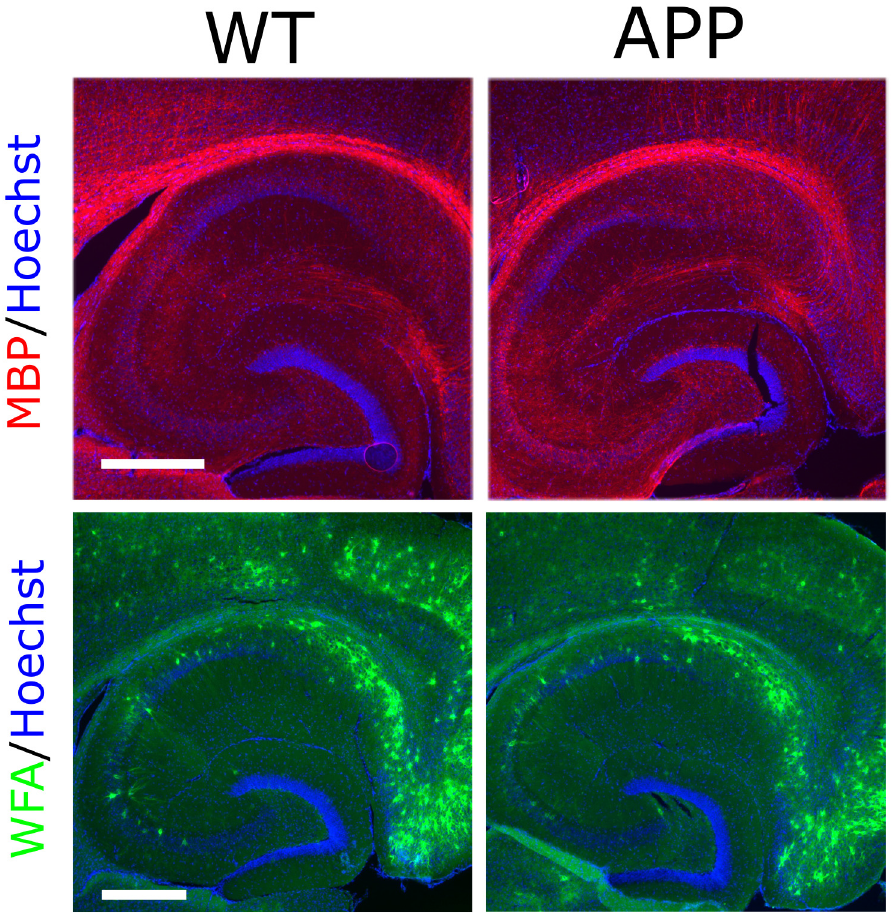
(Immuno)histochemical staining of WT and APP mice hippocampus with MBP for myelin, Hoechst for cell nuclei and WFA for PNNs. Scale bars are 500 μm.

## References

[1] K. Franze, J. Guck, The biophysics of neuronal growth, Reports on Progress in Physics 73 (2010) 094601.

[2] S. P. Lacour, G. Courtine, J. Guck, Materials and technologies for soft implantable neuroprostheses, Nature Reviews Materials 1 (2016) 1–14. Number: 10 Publisher: Nature Publishing Group.

[3] P. Moshayedi, G. Ng, J. C. F. Kwok, G. S. H. Yeo,C. E. Bryant, J. W. Fawcett, K. Franze, J. Guck, The relationship between glial cell mechanosensitivity and foreign body reactions in the central nervous system, Biomaterials 35 (2014) 3919–3925.

[4] T. C, M. C, F. D, V. C, N. A, Glial cell mechanosensitivity is reversed by adhesion cues (2019).

[5] K. Franze, The mechanical control of nervous system development, Development (Cambridge, England) 140 (2013) 3069–3077.

[6] D. E. Koser, A. J. Thompson, S. K. Foster, A. Dwivedy, E. K. Pillai, G. K. Sheridan, H. Svoboda, M. Viana,L. da F. Costa, J. Guck, C. E. Holt, K. Franze, Mechano-sensing is critical for axon growth in the developing brain, Nature neuroscience 19 (2016) 1592–1598.

[7] P. C. Georges, W. J. Miller, D. F. Meaney, E. S. Sawyer,P. A. Janmey, Matrices with compliance comparable to that of brain tissue select neuronal over glial growth in mixed cortical cultures, Biophysical Journal 90 (2006) 3012–3018.

[8] D. Eberle, G. Fodelianaki, T. Kurth, A. Jagielska,S. Möollmert, E. Ulbricht, K. Wagner, A. V. Tauben-berger, N. Traöber, J.-C. Escolano, R. Franklin, K. J. V. Vliet, J. Guck, Acute but not inherited demyelination in mouse models leads to brain tissue stiffness changes, bioRxiv (2018) 449603.

[9] K.-J. Streitberger, I. Sack, D. Krefting, C. Pfuöller,J. Braun, F. Paul, J. Wuerfel, Brain Viscoelasticity Alteration in Chronic-Progressive Multiple Sclerosis, PLoS ONE 7 (2012).

[10] M. M. Urbanski, M. B. Brendel, C. V. Melendez-Vasquez, Acute and chronic demyelinated CNS lesions exhibit opposite elastic properties, Scientific Reports 9 (2019) 999. Number: 1 Publisher: Nature Publishing Group.

[11] L. M. Osborn, W. Kamphuis, W. J. Wadman, E. M. Hol, Astrogliosis: An integral player in the pathogenesis of Alzheimer’s disease, Progress in Neurobiology 144 (2016) 121–141.

[12] M. Fakhoury, Microglia and Astrocytes in Alzheimer’s Disease: Implications for Therapy, Current Neuropharmacology 16 (2018) 508–518.

[13] H. Sasaguri, P. Nilsson, S. Hashimoto, K. Nagata, T. Saito, B. De Strooper, J. Hardy, R. Vassar, B. Winblad, T. C. Saido, APP mouse models for Alzheimer’s disease preclinical studies, The EMBO Journal 36 (2017) 2473–2487.

[14] M. C. Murphy, D. T. Jones, C. R. Jack, K. J. Glaser,M. L. Senjem, A. Manduca, J. P. Felmlee, R. E. Carter,R. L. Ehman, J. Huston, Regional brain stiffness changes across the Alzheimer’s disease spectrum, NeuroImage: Clinical 10 (2016) 283–290.

[15] L. V. Hiscox, C. L. Johnson, M. D. J. McGarry, H. Marshall, C. W. Ritchie, E. J. R. van Beek, N. Roberts,J. M. Starr, Mechanical property alterations across the cerebral cortex due to Alzheimer’s disease, Brain Communications 2 (2020). Publisher: Oxford Academic.

[16] M. Levy Nogueira, S. Epelbaum, J.-M. Steyaert,B. Dubois, L. Schwartz, Mechanical stress models of Alzheimer’s disease pathology, Alzheimer’s & Dementia: The Journal of the Alzheimer’s Association 12 (2016) 324–333.

[17] M. C. Murphy, J. Huston, C. R. Jack, K. J. Glaser,A. Manduca, J. P. Felmlee, R. L. Ehman, Decreased brain stiffness in Alzheimer’s disease determined by magnetic resonance elastography, Journal of magnetic resonance imaging: JMRI 34 (2011) 494–498.

[18] L. M. Gerischer, A. Fehlner, T. Koöbe, K. Prehn, D. Antonenko, U. Grittner, J. Braun, I. Sack, A. Flöoel, Combining viscoelasticity, diffusivity and volume of the hippocampus for the diagnosis of Alzheimer’s disease based on magnetic resonance imaging, NeuroImage: Clinical 18 (2018) 485–493.

[19] n. Kihan Park, G. E. Lonsberry, M. Gearing, A. I. Levey,J. P. Desai, Viscoelastic Properties of Human Autopsy Brain Tissues as Biomarkers for Alzheimer’s Diseases, IEEE transactions on bio-medical engineering 66 (2019) 1705–1713.

[20] G. D. Mahumane, P. Kumar, L. C. d. Toit, Y. E. Choonara, V. Pillay, 3D scaffolds for brain tissue regeneration: architectural challenges, Biomaterials Science 6 (2018) 2812–2837. Publisher: The Royal Society of Chemistry.

[21] M. J. Menal, I. Jorba, M. Torres, J.M. Montserrat,D. Gozal, A. Colell, G. Pinñol-Ripoll, D. Navajas, I. Almendros, R. Farrèe, Alzheimer’s Disease Mutant Mice Exhibit Reduced Brain Tissue Stiffness Compared to Wild-type Mice in both Normoxia and following Intermittent Hypoxia Mimicking Sleep Apnea, Frontiers in Neurology 9 (2018). Publisher: Frontiers.

[22] W. Zhao, W. Cui, S. Xu, L.-Z. Cheong, C. Shen, Examination of Alzheimer’s disease by a combination of electrostatic force and mechanical measurement, Journal of Microscopy 275 (2019) 66–72.

[23] N. Antonovaite, S. V. Beekmans, E. M. Hol, W. J. Wadman, D. Iannuzzi, Regional variations in stiffness in live mouse brain tissue determined by depth-controlled indentation mapping, Scientific Reports 8 (2018).

[24] W. Kamphuis, C. Mamber, M. Moeton, L. Kooijman,J. A. Sluijs, A. H. P. Jansen, M. Verveer, L. R. de Groot,V. D. Smith, S. Rangarajan, J. J. Rodrèıguez, M. Orre,E. M. Hol, GFAP isoforms in adult mouse brain with a focus on neurogenic astrocytes and reactive astrogliosis in mouse models of Alzheimer disease, PloS One 7 (2012) e42823.

[25] D. C. Lin, D. I. Shreiber, E. K. Dimitriadis, F. Horkay, Spherical indentation of soft matter beyond the Hertzian regime: numerical and experimental validation of hyperelastic models, Biomechanics and modeling in mechanobiology 8 (2009) 345–358.

[26] E. G. Herbert, W. C. Oliver, G. M. Pharr, Nanoindentation and the dynamic characterization of viscoelastic solids, Journal of Physics D: Applied Physics 41 (2008) 074021.

[27] Ueber die Beruöhrung fester elastischer Koörper., Journal fuör die reine und angewandte Mathematik (Crelle’s Journal) 1882 (2009) 156–171.

[28] N. Antonovaite, L. A. Hulshof, E. M. Hol, W. J. Wadman, D. Iannuzzi, Viscoelastic mapping of mouse brain tissue: Relation to structure and age, Journal of the Mechanical Behavior of Biomedical Materials (2020) 104159.

[29] E. Galea, W. Morrison, E. Hudry, M. Arbel-Ornath,B. J. Bacskai, T. Gèomez-Isla, H. E. Stanley, B. T. Hy-man, Topological analyses in APP/PS1 mice reveal that astrocytes do not migrate to amyloid-beta plaques, Proceedings of the National Academy of Sciences 112 (2015) 15556–15561. Publisher: National Academy of Sciences Section: Physical Sciences.

[30] M. J. Vèegh, C. M. Heldring, W. Kamphuis, S. Hijazi,A. J. Timmerman, K. W. Li, P. van Nierop, H. D. Mansvelder, E. M. Hol, A. B. Smit, R. E. van Kesteren, Reducing hippocampal extracellular matrix reverses early memory deficits in a mouse model of Alzheimer’s disease, Acta Neuropathologica Communications 2 (2014) 76.

[31] W. Kamphuis, J. Middeldorp, L. Kooijman, J. A. Sluijs, E.-J. Kooi, M. Moeton, M. Freriks, M. R. Mizee, E. M. Hol, Glial fibrillary acidic protein isoform expression in plaque related astrogliosis in Alzheimer’s disease, Neurobiology of Aging 35 (2014) 492–510.

[32] M. C. Murphy, G. L. Curran, K. J. Glaser, P. J. Rossman,J. Huston, J. F. Poduslo, C. R. Jack, J. P. Felmlee, R. L. Ehman, Magnetic resonance elastography of the brain in a mouse model of Alzheimer’s disease: initial results, Magnetic Resonance Imaging 30 (2012) 535–539.

[33] T. Munder, A. Pfeffer, S. Schreyer, J. Guo, J. Braun,I. Sack, B. Steiner, C. Klein, MR elastography detection of early viscoelastic response of the murine hippocampus to amyloid beta accumulation and neuronal cell loss due to Alzheimer’s disease, Journal of Magnetic Resonance Imaging 47 (2018) 105–114.

[34] G. Rosso, I. Liashkovich, V. Shahin, In Situ Investigation of Interrelationships Between Morphology and Biomechanics of Endothelial and Glial Cells and their Nuclei, Advanced Science 6 (2019) 1801638.

[35] J. R. Staunton, B. L. Doss, S. Lindsay, R. Ros, Correlating confocal microscopy and atomic force indentation reveals metastatic cancer cells stiffen during invasion into collagen I matrices, Scientific Reports 6 (2016) 19686. Number: 1 Publisher: Nature Publishing Group.

